# Neutralization of Omicron BA.1, BA.2, and BA.3 SARS-CoV-2 by 3 doses of BNT162b2 vaccine

**DOI:** 10.1101/2022.03.24.485633

**Authors:** Chaitanya Kurhade, Jing Zou, Hongjie Xia, Hui Cai, Qi Yang, Mark Cutler, David Cooper, Alexander Muik, Kathrin U. Jansen, Xuping Xie, Kena A. Swanson, Pei-Yong Shi

**Author notes:** Corresponding authors: X.X., K.A.S., P.-Y.S. H.X. and J.Z. contributed equally to this study. Lead contact: P.-Y.S.

## Abstract

The newly emerged Omicron SARS-CoV-2 has 3 distinct sublineages: BA.1, BA.2, and BA.3. BA.1 accounts for the initial surge and is being replaced by BA.2, whereas BA.3 is at a low prevalence at this time. Here we report the neutralization of BNT162b2-vaccinated sera (collected at 1 month after dose 3) against the three Omicron sublineages. To facilitate the neutralization testing, we engineered the complete BA.1, BA.2, or BA.3 spike into an mNeonGreen USA-WA1/2020 SRAS-CoV-2. All BNT162b2-vaccinated sera neutralized USA-WA1/2020, BA.1-, BA.2-, and BA.3-spike SARS-CoV-2s with titers of >20; the neutralization geometric mean titers (GMTs) against the four viruses were 1211, 336, 300, and 190, respectively. Thus, the BA.1-, BA.2-, and BA.3-spike SARS-CoV-2s were 3.6-, 4.0-, and 6.4-fold less efficiently neutralized than the USA-WA1/2020, respectively. Our data have implications in vaccine strategy and understanding the biology of Omicron sublineages.

## Main text

The severe acute respiratory syndrome coronavirus 2 (SARS-CoV-2) Omicron variant has recently emerged as the fifth variant of concern (VOC) after the previous Alpha, Beta, Gamma, and Delta VOCs. The Omicron variant includes 3 sublineages: BA.1, BA.2, and BA.3. After initial identification in South Africa in November 2021, the Omicron sublineage BA.1, and its derivative BA.1.1 (containing an extra spike R346K substitution), became dominant worldwide. Subsequently, sublineage BA.2 sharply increased its prevalence in many countries, including Demark, the Philippines, South Africa, and Belgium. In the United States, the prevalence of BA.2 increased from 0.4% on January 22 to 23.1% on March 12, 2022 (https://covid.cdc.gov/covid-data-tracker/#variant-proportions). Studies have also shown that BA.2 may be approximately 30% more transmissible than BA.1,^1^ but it does not appear to cause more severe disease.^2^ Compared with the BA.1 and BA.2 sublineages, the prevalence of BA.3 has remained low, with a total of 613 BA.3 sequences in the GISAID database (https://www.gisaid.org/) as of March 12, 2022. Recent studies showed that BA.1 evades vaccine- and non-Omicron infection-elicited neutralization,^3,4^ which, together with increased transmissibility, may partially account for the variant replacement from the previous Delta to the current Omicron. Since BA.1, BA2, and BA.3 have distinct sets of mutations in their spike glycoproteins (**Fig. S1a**), laboratory results are urgently needed to evaluate the susceptibility of the three Omicron sublineages to vaccine-elicited neutralization.

BNT162b2 is an mRNA vaccine that encodes a stabilized prefusion full-length spike glycoprotein from the original SARS-CoV-2 Wuhan-Hu-1 isolate. BNT162b2 mRNA has been approved for vaccination of people ≥16-year-old and has also been authorized under emergency use provision for immunization of children 5-to 15-year-old by the US Food and Drug Administration. To determine the susceptibility of Omicron sublineages to BNT162b2-elicited neutralization, we engineered the complete BA.1 (GISAID EPI_ISL_6640916), BA.2 (GISAID EPI_ISL_6795834.2), or BA.3 (GISAID EPI_ISL_7605591) spike into the mNeonGreen (mNG) reporter USA-WA1/2020, an SARS-CoV-2 strain isolated in January 2020 (**Fig. S1a**). The mNG gene was inserted into the open-reading-frame-7 (ORF7) of the viral genome to enable the development of a fluorescent focus reduction neutralization test (FFRNT).^4^ Our historical data set indicate that the high-throughput FFRNT produces comparable neutralization results as the gold standard plaque-reduction neutralization test (PRNT; using SARS-CoV-2 without the mNG reporter) when analyzing BNT162b2-vaccinated human sera (**Fig. S2**). Thus, FFRNT can reliably be used to measure antibody neutralization.

Characterization of the recombinant BA.1-, BA.2-, and BA.3-spike mNG SARS-CoV-2s showed that all viruses produced infectious titers of >10^6^ focus-forming units per milliliter (FFU/ml), similar to the wild-type USA-WA1/2020 mNG virus. Although the recombinant viruses formed different sizes of fluorescent foci in the order of wild-type USA-WA1/2020 > BS.2-spike ≈ BA.3-spike > BA.1-spike mNG SARS-CoV-2 (**Fig. S1b**), all viruses showed equivalent viral RNA genome/FFU ratios when analyzed on Vero E6 cells (**Fig. S1c**), suggesting equivalent specific infectivities of the viral stocks. All recombinant viruses were sequenced to ensure no undesired mutations.

Using a panel of human sera collected at 1 month post dose 3 (PD3) of BNT162b2 vaccine^3^ (**Fig. S3**), we determined the 50% fluorescent focus-reduction neutralization titers (FFRNT_50_) against recombinant USA-WA1/2020 and Omicron sublineage-spike mNG SARS-CoV-2s. We chose the PD3 sera, rather than post dose 2 sera, because (i) two doses of BNT162b2 did not elicit robust neutralization against Omicron BA.1^3^ and (ii) many individuals have already received 3 doses of BNT162b2. The PD3 sera neutralized USA-WA1/2020, BA.1-, BA.2-, and BA.3-spike mNG viruses with geometric mean titers (GMTs) of 1211, 336, 300, and 190, respectively (**Fig. 1**). Although all PD3 sera neutralized recombinant viruses with titers of >20, the neutralizing GMTs against BA.1-, BA.2-, and BA.3-spike mNG viruses were 3.6-, 4.0-, and 6.4-fold lower than the GMT against the mNG USA-WA1/2020, respectively (**Fig. 1**). The results support two conclusions. First, BA.1 and BA.2 spikes evade the neutralization of PD3 sera to comparable degrees. Similar reduced neutralization of BA.1 and BA.2 sublineages were reported;^5,6^ however, the reduction folds varied among studies, likely due to different serum specimens and neutralization assay protocols. These results may also imply that the observed increasing prevalence of BA.2 over BA.1 among circulating strains was not driven by the difference in antibody neutralization after vaccination, but by other factors, such as differences in viral replication^7^ and transmission,^8^ or other potential immune evasion mechanisms. Second, BA.3 spike evades the BNT162b2-elicited neutralization more efficiently than BA.1 and BA.2 spikes. If the viral replication of BA.3 is not attenuated (which remains to be determined), the BA.3 sublineage may have the potential to expand and elongate the current Omicron surges. Thus, we should closely monitor the prevalence of BA.3 during surveillance.

**Figure 1.**
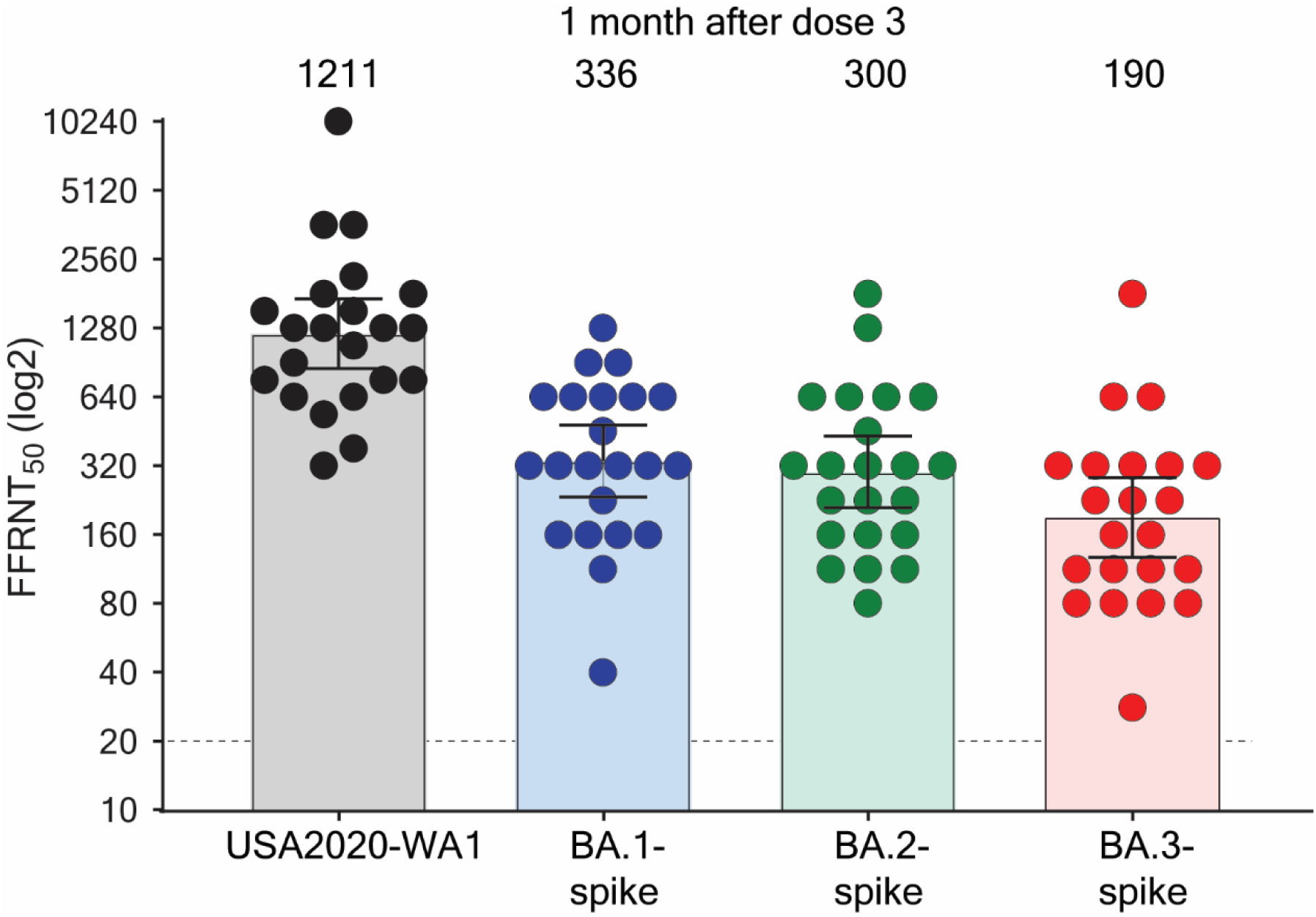
Serum neutralization of Omicron BA.1-, BA.2-, and BA.3-spike mNG SARS-CoV-2s and USA-WA1/2020 after three doses of BNT162b2. A panel of 22 human sera collected at 1 month after dose 3 of BNT162b2 vaccine were tested for the 50% fluorescent focus-reduction neutralization titers (FFRNT_50_) against recombinant USA-WA1/2020, Omicron BA.1-, BA.2-, and BA.3-spike mNG SARS-CoV-2s. The serum collection scheme is described in **Figure S3** as previously reported.^19^ The BA.1-, BA.2-, and BA.3-spike mNG SARS-CoV-2s were produced by engineering the complete Omicron spike genes into the mNG USA-WA1/2020.^15^ Each data point represents the geometric mean FFRNT_50_ obtained with a serum specimen against the indicated virus, as detailed in **Tables S1**. The neutralization titers for BA.1-, BA.2-, and BA.3-spike mNG SARS-CoV-2s were determined in duplicate assays; the FFRNT_50_s for USA-WA1/2020 mNG SARS-CoV-2 were determined in two independent experiments, each with duplicate assays; the geometric means were presented. The bar heights and the numbers above indicate geometric mean titers. The whiskers indicate 95% confidence intervals. The dotted line indicates the limit of detection of FFRNT_50_. Statistical analysis was performed with the use of the Wilcoxon matched-pairs signed-rank test. The statistical significances of the differences between geometric mean titers against the USA-WA1/2020 and Omicron BA.1-, BA.2-, or BA.3-spike SARS-CoV-2 are all p < 0.0001.

Despite the reduced neutralization of Omicron sublineages, other immune effectors, such as T cells and non-neutralizing antibodies that mediate antibody-dependent cytotoxicity, also contribute to the protection against severe COVID-19. The majority of T cell epitopes after vaccination or natural infection are preserved against Omicron spikes.^9^ Indeed, 3 doses of BNT162b2 has conferred efficacy against Omicron disease; however, recent real world effectiveness data showed a waning of protection against symptomatic infection caused by Omicron. Waning of protection was more modest against severe disease caused by Omicron, with overall efficacy remaining high 3 to 6 months after dose 3.^10-14^ Studies are underway to determine the extended durability of the PD3 neutralization against Omicron sublineages and future variants. These laboratory investigations, together with real world vaccine effectiveness data, will continue to guide 4^th^ dose vaccine strategy for optimal breadth and duration of protection.

## Methods

### Construction and characterization of recombinant Omicron sublineage spike mNG SARS-CoV-2s

Recombinant Omicron BA.1-, BA.2-, and BA.3-spike mNG SARS-CoV-2s were constructed by engineering the complete spike gene from Omicron sublineages into an infectious cDNA clone of mNG USA-WA1/2020^15^ (**Figure S1a**). All spike mutations, deletions, and insertions were introduced into the infectious cDNA clone of mNG USA-WA1/2020 using PCR-based mutagenesis as previously described.^16^ *The BA.1, BA.2, and BA.3 spike sequences were based on GISAID* EPI_ISL_6640916, EPI_ISL_6795834.2, and EPI_ISL_7605591, respectively. The full-length cDNA of viral genome containing the complete Omicron spike was assembled via *in vitro* ligation. The resulting full-length cDNA was used as a template for *in vitro* transcription of genome-length viral RNA. The *in vitro* transcribed viral RNA was electroporated into Vero E6 cells. On day 3 post electroporation, the original viral stock (P0) was harvested from the electroporated cells. The P0 virus was amplified for another round on Vero E6 cells to produce the P1 stock for neutralization testing. The infectious titer of the P1 virus was quantified by fluorescent focus assay on Vero E6 cells (**Figure S1b**). We sequenced the complete spike gene of the P1 virus to ensure no undesired mutations. Only the P1 viruses were used for neutralization test. The protocols for the mutagenesis of mNG SARS-CoV-2 and virus production were reported previously.^17^ To determine the specific infectivity, we quantified the P1 virus stocks for their fluorescent focus units (FFU) and genomic RNA contents by fluorescent focus assay on Vero E6 cells and RT-qPCR, respectively. The methods for fluorescent focus assay and RT-qPCR were reported previously.^4,18^ The specific infectivity of each virus was measured by the genomic RNA-to-FFU ratios (genome/FFU).

### Fluorescent focus reduction neutralization test (FFRNT)

Neutralization titers of human sera were measured by FFRNT using the USA-WA1/2020, BA.1-, BA.2-, and BA.3-spike mNG SARS-CoV-2s. The details of FFRNT protocol were reported previously.^4^ Briefly, 2.5×10^4^ Vero E6 cells per well were seeded in 96-well plates (Greiner Bio-one™). The cells were incubated overnight. On the next day, each serum was 2-fold serially diluted in the culture medium with the first dilution of 1:20 (final dilution range of 1:20 to 1:20,480). The diluted serum was incubated with 100-150 FFUs of mNG SARS-CoV-2 at 37 °C for 1 h, after which the serum-virus mixtures were loaded onto the pre-seeded Vero E6 cell monolayer in 96-well plates. After 1 h infection, the inoculum was removed and 100 μl of overlay medium (supplemented with 0.8% methylcellulose) was added to each well. After incubating the plates at 37 °C for 16 h, raw images of mNG foci were acquired using Cytation™ 7 (BioTek) armed with 2.5× FL Zeiss objective with wide field of view and processed using the software settings (GFP [469,525] threshold 4000, object selection size 50-1000 μm). The foci in each well were counted and normalized to the non-serum-treated controls to calculate the relative infectivities. The FFRNT_50_ value was defined as the minimal serum dilution that suppressed >50% of fluorescent foci. The neutralization titer of each serum was determined in duplicate assays, and the geometric mean was taken. **Tables S1** summarizes the FFRNT_50_ results.

## Acknowledgements

We thank the Pfizer-BioNTech clinical trial C4591001 and NCT04368728 participants, from whom the post-immunization human sera were obtained. We thank the many colleagues at Pfizer and BioNTech who developed and produced the BNT162b2 vaccine.

## Author contributions

C.K., K.U.J., X.X., K.A.S., and P.-Y.S. conceived the study. C.K., J.Z., H.X., H.C., Q.Y., M.C., D.C., X.X., and K.A.S. performed the experiments. H.X., J.Z., M.C., D.C., A.M., K.U.J., X.X., K.A.S., and P.-Y.S. analyzed the results. C.K., J.Z., X.X., K.A.S., and P.-Y.S. wrote the manuscript. M.C., D.C., K.U.J., X.X., K.A.S., and P.-Y.S. supervised the project.

