## Supplementary material for "Neutralization of Omicron BA.1, BA.2, and BA.3 SARS-CoV-2 by 3 doses of BNT162b2 vaccine": Methods Supplementary figures 1-3 and Tables 1-2

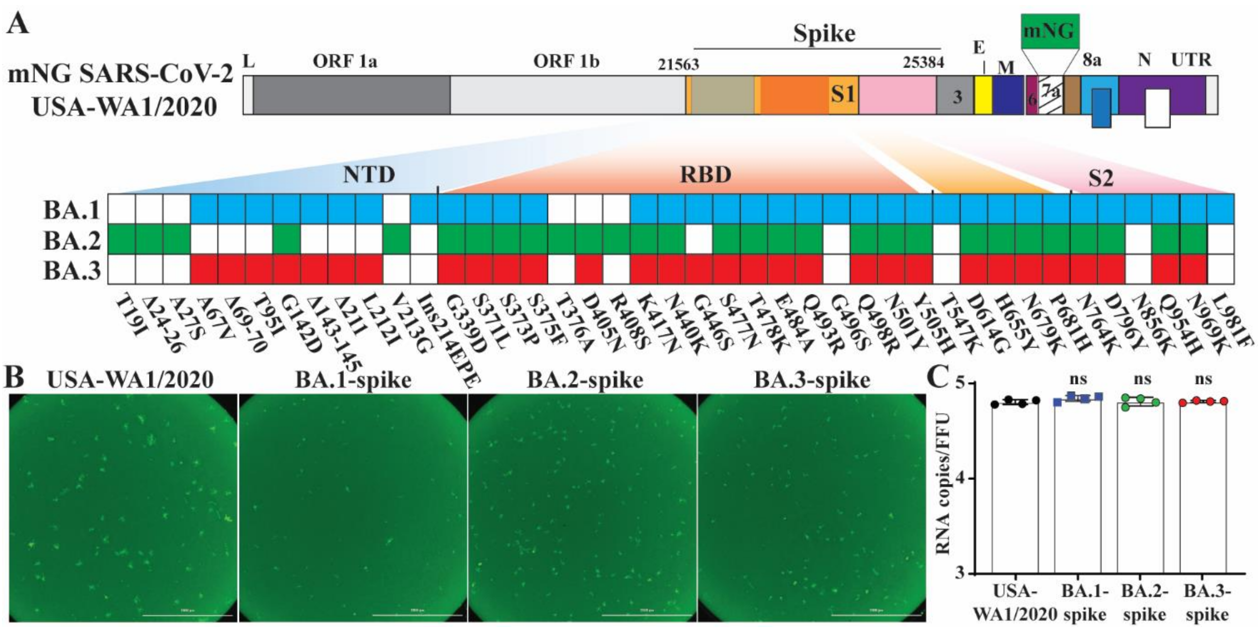


**Figure S1. Construction and characterization of Omicron subvariant BA.1-, BA.2-, and BA.3-spike mNG SARS-CoV-2s.** (A) Omicron BA.1-, BA.2-, and BA.3-spike mNG SARS-CoV-2s. mNG USA-WA1/2020 was used to engineer Omicron subvariant spike SARS-CoV-2s. The mNG reporter gene was engineered at the open-reading-frame-7 (ORF7) of the USA-WA1/2020 genome. Amino acid mutations, deletions (Δ), and insertions (Ins) are indicated for BA.1, BA.2, and BA.3 spikes in comparison with the USA-WA1/2020 spike. L: leader sequence; ORF: open reading frame; NTD: N-terminal domain of S1; RBD: receptor binding domain of S1; S: spike glycoprotein; S1: N-terminal furin cleavage fragment of S; S2: C-terminal furin cleavage fragment of S; E: envelope protein; M: membrane protein; N: nucleoprotein; UTR: untranslated region. (B) Fluorescent foci formed by mNG USA-WA1/2020, BA.1-, BA.2-, and BA.3-spike mNG SARS-CoV-2s. The images were taken at 16 h after Vero E6 cells were infected with the indicated viruses in 96-well plates. (C) Specific infectivity. The specific infectivity of mNG USA-WA1/2020 or Omicron subvariant spike mNG SARS-CoV-2 was determined by the ratio of viral genomic RNA versus fluorescent focus-forming unit ratios (genomes/FFU). Individual recombinant virus P1 stocks were measured for their viral RNA and FFUs by RT-qPCR and fluorescent focus assay on Vero E6 cells, respectively. Dots represent individual biological replicates of 4 different aliquots of viruses. The values in the graph represent the means with 95% confidence intervals. A non-parametric Mann-Whitney test was used to determine significant difference between USA-WA1/2020 and Omicron spike mNG SARS-CoV-2s. n.s., no statistical difference.

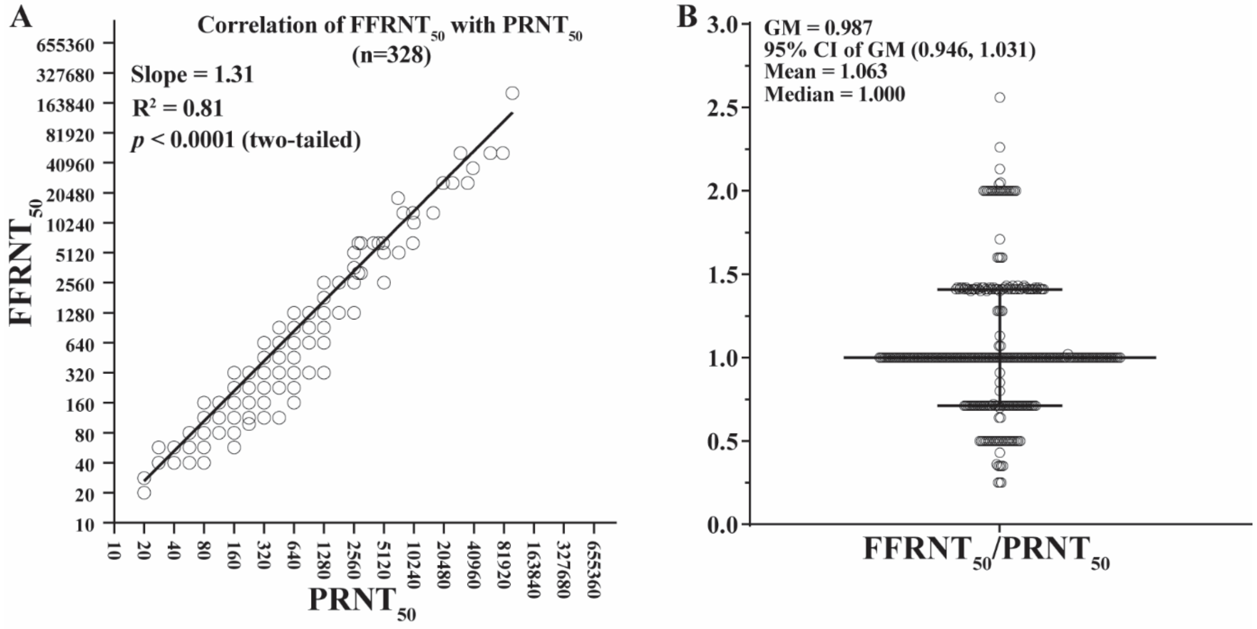


**Figure S2. Correlation of neutralization titers measured by fluorescent focus-reduction neutralization test (FFRNT) and plaque-reduction neutralization test (PRNT).** A historical data set of BNT162b2-vaccinated sera are presented with a total of 328 tests. The neutralization titers of a serum panel were measured using mNG SARS-CoV-2-based FFRNT and conventional SARS-CoV-2-based PRNT methods. The correlation between FFRNT$_{50}$ and PRNT$_{50}$ is plotted in (A). The FFRNT$_{50}$/PRNT$_{50}$ ratios are presented in (B). GM, geometric mean; CI, confidence interval. Error bar shows the 95% CI of GM.

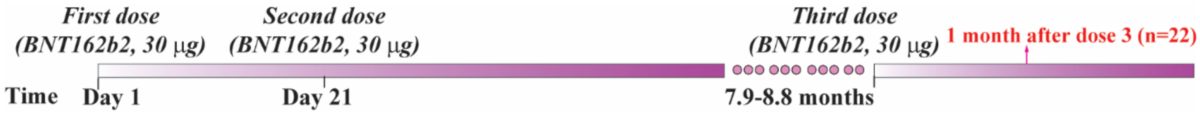


**Figure S3. BNT162b2-vaccinated sera.** A panel of 22 human sera were collected at 1 month post 3 doses of BNT162b2 vaccine. The time intervals between the three doses are indicated. This panel of sera was as recently reported.[1]

### Table S1. $FFRNT_{50}$ values of BNT162b2-vaccinated sera

| Subject ID | Age (Years) | Sex (F/M) | *$FFRNT_{50}$ | | | | | |
|---|---|---|---|---|---|---|---|---|
| | | | USA-WA1/2022 | | | BA.1-spike | BA.2-spike | BA.3-spike |
| | | | [&]Exp1 | [&]Exp2 | GMT | | | |
| 1 | 26 | F | 640 | 905 | 761 | 320 | 320 | 226 |
| 2 | 28 | M | 1280 | 1810 | 1522 | 320 | 320 | 113 |
| 3 | 35 | F | 1280 | 1280 | 1280 | 640 | 453 | 320 |
| 4 | 35 | F | 1280 | 1280 | 1280 | 320 | 226 | 226 |
| 5 | 38 | F | 1280 | 1810 | 1522 | 640 | 640 | 320 |
| 6 | 38 | F | 320 | 320 | 320 | 160 | 160 | 80 |
| 7 | 44 | F | 640 | 453 | 538 | 113 | 113 | 80 |
| 8 | 44 | F | 2560 | 1280 | 1810 | 640 | 640 | 320 |
| 9 | 52 | F | 2560 | 1280 | 1810 | 320 | 320 | 320 |
| 10 | 54 | M | 1280 | 1280 | 1280 | 453 | 320 | 226 |
| 11 | 65 | M | 1280 | 640 | 905 | 226 | 226 | 113 |
| 12 | 65 | M | 5120 | 2560 | 3620 | 640 | 1280 | 320 |
| 13 | 66 | F | 640 | 640 | 640 | 160 | 160 | 80 |
| 14 | 67 | M | 640 | 905 | 761 | 320 | 160 | 113 |
| 15 | 68 | M | 640 | 905 | 761 | 160 | 113 | 113 |
| 16 | 68 | F | 5120 | 2560 | 3620 | 905 | 640 | 640 |
| 17 | 68 | F | 640 | 640 | 640 | 160 | 113 | 80 |
| 18 | 69 | F | 1280 | 905 | 1076 | 320 | 226 | 160 |
| 19 | 69 | F | 2560 | 1810 | 2153 | 905 | 640 | 640 |
| 20 | 70 | M | 10240 | 10240 | 10240 | 1280 | 1810 | 1810 |
| 21 | 73 | M | 320 | 453 | 381 | 40 | 80 | 28 |
| 22 | 74 | F | 1280 | 1280 | 1280 | 640 | 320 | 160 |
| [†]GMT | -- | -- | 1280 | 1146 | 1211 | 336 | 300 | 190 |
| [^]95% CI | -- | -- | 866-1893 | 827-1589 | 854-1718 | 234-482 | 210-431 | 127-284 |

*Individual $FFRNT_{50}$ value is the geometric mean of duplicate plaque assay results.
[&]Two independent experiments were performed for mNG USA-WA1/2022 because BA.1-, BA.2-, and BA.3-spike SARS-CoV-2s were tested in two experiments.
[#]The sera were collected at 1 month after dose 3 of BNT162b2 vaccine, as recently reported.[1]
[†]Geometric mean neutralizing titers.
[^]95% confidence interval (95% CI) for the GMT.
